# MinSNPs: an R package for derivation of resolution-optimised SNP sets from microbial genomic data

**DOI:** 10.1101/2022.07.27.501805

**Authors:** Kian Soon Hoon, Deborah C Holt, Sarah Auburn, Peter Shaw, Philip M. Giffard

**Affiliations:** Division of Global and Tropical Health, Menzies School of Health Research, Charles Darwin University, Darwin, Northern Territory, Australia; College of Health and Human Sciences, Charles Darwin University, Darwin, Northern Territory, Australia; Mahidol-Oxford Tropical Medicine Research Unit, Mahidol University, Bangkok, Thailand; Centre for Tropical Medicine and Global Health, Nuffield Department of Medicine, University of Oxford, Oxford, United Kingdom; Oujiang Laboratory, Wenzhou, Zhejiang, China

**Keywords:** SNP, Bacteria, Plasmodium, Staphylococcus, resolution optimised, genotyping, surveillance, SNP matrix, SNP set derivation, genetic epidemiology

## Abstract

2.

Here we present the R package - MinSNPs. This is designed to assemble resolution optimised sets of single nucleotide polymorphisms (SNPs) from alignments such as genome wide orthologous SNP matrices. We also demonstrate a pipeline for assembling such matrices from multiple bio-projects, so as to facilitate SNP set derivation from globally representative data sets. MinSNPs can derive sets of SNPs optimised for discriminating any user-defined combination of sequences from all others. Alternatively, SNP sets may be optimised to discriminate all from all, i.e., to maximise diversity. MinSNPs encompasses functions that facilitate rapid and flexible SNP mining, and clear and comprehensive presentation of the results. The MinSNPs running time scales in a linear fashion with input data volume, and the numbers of SNPs and SNPs sets specified in the output. MinSNPs was tested using a previously reported orthologous SNP matrix of *Staphylococcus aureus*. and an orthologous SNP matrix of 3,279 genomes with 164,335 SNPs assembled from four *S. aureus* short read genomic data sets. MinSNPs demonstrated efficacy in deriving discriminatory SNP sets for potential surveillance targets and in identifying SNP sets optimised to discriminate isolates from different clonal complexes (CC). MinSNPs was also tested with a large *Plasmodium vivax* orthologous SNP matrix. A set of five SNPs was derived that reliably indicated the country of origin within 3 south-east Asian countries. In summary, we report the capacity to assemble comprehensive SNP matrices that effectively capture microbial genomic diversity, and to rapidly and flexibly mine these entities for optimised surveillance marker sets.

**Impact statement:** We present the R package “MinSNPs”. This derives resolution optimised SNP sets from datasets of genome sequence variation. Such SNP sets can underpin targeted genetic analysis for high throughput surveillance of microbial variants of public health concern. MinSNPs supports considerable flexibility in search methods. The package allows non-specialist bioinformaticians to easily and quickly convert global scale data of intra-specific genomic variation into SNP sets precisely and efficiently directed towards many microbial genetic analysis tasks.

**Data summary:**

1. The source code for minSNPs is available from GitHub under MIT Licence (URLs – https://github.com/ludwigHoon/minSNPs and mirrored in https://cran.r-project.org/package=minSNPs)
2. *Staphylococcus aureus* (STARRS data set) Orthologous SNP Matrix; (URL - https://doi.org/10.1371/journal.pone.0245790.s005)
3. *Plasmodium vivax* data set (VCF file); (URL - https://www.malariagen.net/resource/24)
4. *Staphylococcus aureus* short read sequences (fastq) from bioprojects: PRJEB40888 (or STARRS)(https://www.ncbi.nlm.nih.gov/bioproject/PRJEB40888), PRJEB3174 (https://www.ncbi.nlm.nih.gov/bioproject/PRJEB3174), PRJEB32286 (https://www.ncbi.nlm.nih.gov/bioproject/PRJEB32286), and PRJNA400143 (https://www.ncbi.nlm.nih.gov/bioproject/PRJNA400143)

## 5. Introduction

The extremely large-scale accumulation of microbial whole genome sequence information provides a potent resource for the design of targeted genetic analysis procedures. While whole genome analysis is now widely applied directly to public health, clinical, and research microbiology, targeted genetic analyses may be complementary to whole genome analysis for purposes such as high-volume, low-cost surveillance, analysis of primary specimens, and/or analyses performed outside the laboratory environment. Several research groups have recently developed SNP-based genotyping approaches, e.g., to investigate *Mycobacterium species* (1, 2), attribute host for *Chlamydia psittaci* (3) and *Campylobacter coli (4)*, distinguish *Rickettsia typhi* from different continents (5), identify *Escherichia coli* of specific serotype (6), and track the spread of drug resistance in *Plasmodium falciparum* infections (7).

Here we report the R package “MinSNPs”. This package is designed to derive sets of polymorphisms from biological sequence alignment data on the basis of high combinatorial discriminatory power. The envisioned application is the derivation of high-resolution sets of single nucleotide polymorphisms (SNPs) from DNA sequence alignments or orthologous SNP matrices. MinSNPs encompasses much of the functionality of the previously reported “Minimum SNPs” Java-based bioinformatics application (8, 9). Minimum SNPs was used to develop a number of SNP-based bacterial genotyping methods e.g., (10–14). MinSNPs is a new package, written in R, with distinct code from Minimum SNPs. The reasons for re-development were improvement of flexibility, error handling, and output formats.

Here we describe MinSNPs and demonstrate functionality using comparative genome data from *Staphylococcus aureus* and *Plasmodium vivax*. We also demonstrate a pipeline to generate MinSNPs input files from multiple short read data sets to facilitate the analysis of data from multiple studies.

## 6. Theory & implementation

The input format is a single sequence alignment in FASTA format. All symbols are recognised so that the program will derive sets of polymorphic positions from any file in a FASTA format alignment, irrespective of the symbols in the sequences. However, symbols that are not G, A, T or C can optionally trigger the exclusion of the relevant alignment positions from analysis. Our focus has largely been on the analysis of genome-wide orthologous SNP matrices.

The output of MinSNPs is set(s) of polymorphic positions in the alignment. SNP sets are assembled iteratively, on the basis of maximised combinatorial resolving power. SNP 1 is the single SNP with the highest resolving power, SNP 2 is the SNP with the highest resolving power in combination with SNP 1 etc. Where more than one SNP confers the same increase in resolving power, the SNP nearest to position 1 will be added to the set.

There are two user-selectable algorithms for measuring resolving power.

1. **% mode.** The resolving power is the percentage of sequences in the alignment that are not discriminated from the user-selected sequence(s) (the group of interest). The SNP sets are constrained to 100% sensitivity. The first SNP identified is the 100% sensitive SNP with maximum possible specificity, while subsequent SNPs are selected on the basis of the maximum possible increase in specificity in combination with the previously selected SNP(s). All alignment positions that are variable within the group of interest can optionally be excluded from the analysis. We suggest that, where possible, the group of interest be composed of >1 sequence to avoid the identification of spurious SNPs arising from sequencing errors.
2. ***D* mode**. The resolving power is the power to discriminate “all from all”, as measured by the Simpsons index of diversity (*D*). In this context, *D* is the probability that any two sequences in the alignment will be discriminated from each other by the SNP set, as calculated by 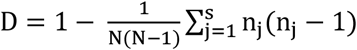, where N is the number of sequences, s is the number of classes defined by the SNPs, and nj is the number of sequences defined by the class j (15).

MinSNPs encompasses functions that support flexibility of analyses and transparency of outputs. The user specifies the size and number of the SNP sets that constitute the output. When multiple SNP sets are requested, MinSNPs identifies alternative SNP sets that are all resolution optimised, with the constraint that the sets must differ from each other at least in the first SNP.

The user can force the program to include or exclude any alignment position(s) in/from the SNP set. Where positions are included, new SNPs are identified based on resolving power in combination with the included positions. This facilitates rapid modification of SNP sets.

MinSNPs can identify alignment positions where at least one sequence has a non-standard DNA symbol, and these positions are optionally excluded from analysis. Indels (dashes) default to being regarded as symbols equivalent to other symbols. Alternatively, the user can specify that indels trigger the exclusion of the relevant alignment positions from the analysis. There is also an optional function to exclude positions with SNPs with >2 alleles.

MinSNPs provides a cumulative increase in resolving power as the sets are built, and the tabulated information indexing the sequences in the alignment as defined by each allelic profile. For % mode analyses, this is within a “group of interest or non-group of interest” framework. The outputs are presented in the R console and optionally outputted to a tab-delimited format file. A facile method to fully define the informative power of a SNP set derived by % analysis is to force the inclusion of the SNPs into a *D* analysis, in which the user-defined SNP set size equals the number of included SNPs, i.e., no additional SNPs are derived. This will reveal how the sequences assort in relation to allelic profiles of the “forced included” SNPs. Alternatively, this can be done in reverse to assess the performance of a *D* maximised SNP set to detect user-defined subsets of sequences with 100% sensitivity. These functions provide considerable flexibility regarding the exploration of SNP sets.

### 6.1 Demonstration of MinSNPs functionality

To explore the potential utility of MinSNPs, we:

1. Determined the relationship between input alignment dimensions and the number and size of output SNP sets, with running time;
2. Generated SNP sets of potential relevance to surveillance from orthologous SNP matrices derived from genomic epidemiology studies in *Staphylococcus aureus* and *Plasmodium vivax;*
3. Explored the properties of SNPs identified with MinSNPs with respect to genome position and relationship with coding sequences and;
4. Developed a pipeline to generate single orthologous SNP matrices from multiple short-read data sets. This can support the analysis of large-scale comparative genome data using MinSNPs.

#### 6.1.1 Run-time Determinations

The relationships between the analysis time and dimensions of the input alignment, the number of SNPs in the output SNP set, and the number of SNP sets in the output were determined. The relationship was linear with respect to all three parameters. Examples of running time are shown in **Error! Reference source not found.**. It was also shown that running MinSNPs using multiple cores improves its performance. Complete data and code are shown in Supplementary Information 1 (https://figshare.com/s/696f696c232404f18a36). The faster run-time on a laptop as compared to a high-performance cluster (HPC) was due to the simpler architecture of the machine; we note that when the dimension of the alignments increases, the HPC’s performance improves. So, given a higher number of cores and increased memory available, a HPC can easily outperform a laptop.

#### 6.1.2 Derivation of SNP sets from a *Staphylococcus aureus* orthologous SNP matrix

To demonstrate MinSNPs’ functionality, we analysed genome-wide orthologous SNP matrices to identify 1. SNP sets diagnostic for a conserved lineage that is a potential surveillance target, 2. SNP sets diagnostic for a broader phylogenetic lineage that encompasses the potential surveillance target, and 3. SNP sets optimised with respect to *D*. For the latter, our interests were in the resolving power (the *D* value), and the concordance of the genotypes defined by the SNP sets with the phylogeny indicated by the orthologous SNP matrix.

We first analysed a previously described orthologous SNP matrix composed of 20,651 SNPs from 162 *S. aureus* isolates, four *Staphylococcus argenteus* isolates, and *S. aureus* Mu50, which was the reference genome for matrix construction (13). The isolates were from the STARRS study, which revealed potential *S. aureus* transmission events involving haemodialysis patients, and potential contacts in the clinical context environment, in the north of the Australian Northern Territory (13).

The STARRS study identified isolates of multilocus sequence typing (MLST) defined ST762 (clonal complex (CC) 1), and were involved in transmission events leading to patient infections. ST762 is vanishingly rare globally but was prevalent in the STARRs study. We therefore used the ST762 lineage identified in the STARRs study as a model for a potential surveillance target. Using MinSNPs in % mode, we determined that 12 SNPs each individually discriminated all the ST762 isolates from other isolates in the study, with 100% sensitivity and specificity (Supplementary Information 2 (https://figshare.com/s/746dd263140963185c53)). A BLAST analysis demonstrated that for each of these SNPs, the alleles present in the ST762 isolates were not present in the public databases, suggesting that these SNPs have generalised ability to discriminate ST762 from the remainder of the *S. aureus* complex (Supplementary Information 3 (https://figshare.com/s/8adc2b14052ccb89dbed)).

The same procedure was used to derive SNP sets that discriminate the CC1 (ST1 and ST762) STARRS isolates from the other isolates. It was found that there were 119 SNPs that each individually provided 100% sensitivity and specificity (Supplementary Information 2). Similar to SNPs identified for ST762, a BLAST analysis returned 61 specimens from Genbank; out of these 53 are CC1 with 3 false positives belonging to ST425 and 5 specimens untypeable by MLST.

We further used MinSNPs to derive 15 five-member SNP sets with maximised *D*. The *D* values obtained ranged from 0.925 to 0.936, defining 16 to 21 genotypes. Concordance with phylogeny was determined for two SNP sets (set 1 and 11) that were selected on the basis of having no SNPs in common. Both SNP sets discriminate the major lineages defined by the STARRS SNP matrix (**Error! Reference source not found.**, **Error! Reference source not found.**).

#### 6.1.3 Derivation of *Plasmodium vivax SNP sets*

Given challenges associated with the large genome size and high proportions of ‘contaminating’ human DNA, targeted SNP genotyping remains an important approach in *Plasmodium* epidemiological tracking (16–18). MinSNPs was tested with a *P. vivax* orthologous SNP matrix encompassing 259 isolates and 527,107 SNPs (19). The matrix encompassed heterozygote positions (read as nucleotide ambiguities in MinSNPs) that enabled us to develop strategies to accommodate this feature, which is common in polyclonal infections.

The data were generated from isolates collected from Malaysia, Thailand, and Indonesia, as part of a study to identify changes in the *P. vivax* population as Sabah (Malaysia) approaches the elimination of vivax malaria (19). In 183,509 of the SNPs, a nucleotide ambiguity code (where calls were heterozygote) was assigned to at least one of these isolates.

As previously described, a subset of 26 specimens from Malaysia were near identical. These were denoted “K2” strains reflecting isolates that were potentially undergoing clonal expansion (19). We regarded these as a model surveillance target. SNPs that discriminated the K2 lineage were identified with MinSNPs in % mode, with all the K2 specimens defined as the group of interest. All 183,509 positions where any of the sequences had an ambiguity code were excluded from the analysis. The resulting analysis of 343,598 SNPs yielded 124 SNPs that each individually discriminated the K2 lineage from all the other isolates in the matrix (Supplementary Information 4 https://figshare.com/s/db47a069aab93f3c615c)). Any of these 124 SNPs could potentially form the basis of a K2 surveillance tool protocol, and using more than one of these SNPs may provide useful redundancy to avoid false negatives due to undiscovered sequence diversity.

Next, SNPs that discriminated all Malaysian specimens from all other specimens were derived. To streamline the analysis, only one K2 specimen was included. Also, three specimens that were obtained in Malaysia but were likely to be imported from other regions based on their genomic clustering patterns (PY0045-C, PY0004-C and PY0120-C) were omitted from the group of interest. Initially, we confined the analysis to the 343,598 SNPs that do not encompass any ambiguity codes. This was not successful. The maximum % obtained from five SNPs was 0.265, meaning that 73.5% of the non-Malaysian specimens were not discriminated from the Malaysian specimens (Supplementary Information 4). A different protocol was then adopted. Prior to MinSNPs analysis, ambiguity codes were transformed into the major allele at that position (Supplementary Information 4 (https://figshare.com/s/db47a069aab93f3c615c)). Fortuitously, in all cases, the major allele was consistent with the ambiguity code. After MinSNPs analysis, the relationship between the allelic profiles and isolate was determined using the untransformed matrix. The untransformed matrix can define allelic profiles that include ambiguity codes. Any specimens that had such an allelic profile, i.e., they had an ambiguity code at a SNP within the SNP set being assessed, were classified as untypeable by that SNP set. Typeability was therefore a criterion we used for assessing SNP sets, although we do note that typeability is likely a function of specimen quality and/or whether the specimen contained a mixture of strains. It is not an inherent property of a pure *P. vivax* clone.

This approach to identifying SNPs that discriminated Malaysian specimens was successful. Two sets of two SNPs were identified, each of which discriminated all Malaysian specimens from all other typable specimens. For one SNP set, 20 specimens (7.72%) were untypeable, and for the other, the number of untypeable specimens was 22 (8.49%). All the Malaysian specimens were typable with both SNP sets. The reason for the superior result from the matrix with ambiguity codes transformed is unclear. However, we note that the MinSNPs’ requirement in % mode that SNP sets provide 100% sensitivity for the group of interest, is a stringent constraint. A false negative defined by a single member of a group of interest disqualifies a position from inclusion in a SNP set. Being able to capture more diversity for the analysis by using the transformation procedure also appears to have been critical. A possible work-around for this constraint on SNP selection is to run separate analyses, each with subsets of the group of interest.

We then used MinSNPs to derive *D* maximised SNP sets from the *P. vivax* alignment. Both the approaches described above for accommodating ambiguity codes were used. Five SNP sets, each comprising five SNPs were derived using each approach. When all the positions that encompassed at least one ambiguity code were excluded from the analysis, the *D* values obtained were 0.751, 0.750, 0.572, and 0.564 (two sets). The most discriminatory SNP set (D = 0.751) was investigated further. It was determined that the matrix defined eight allelic profiles. Although this number of profiles and the *D* value do not indicate high discrimination, there was close concordance between allelic profile and country of origin ((Supplementary Information 4 (https://figshare.com/s/db47a069aab93f3c615c), **Error! Reference source not found.**). Thus, within the context of the diversity defined by the input matrix, five SNPs can accurately reveal *P. vivax* country of origin. When the analysis was repeated with the transformed ambiguity codes, very different results were obtained. The *D* values were between 0.958 to 0.960, which is considerably higher than in the previous experiment. Consistent with this, the SNP sets defined 31-32 allelic profiles. The numbers of specimens defined as untypeable were significant, ranging from 64 to 68 (25%-26% of specimens). The concordance between country of origin was poor. Even with the larger number of allelic profiles, there were numerous instances of specimens from different countries having the same profile. A likely explanation is that positions that encompass ambiguity codes are polymorphic within countries. Such SNPs are more likely to generate ambiguity codes because both alleles may be present in a mixed infection. The exclusion of these positions will enrich for SNPs that separate specimens from different countries and are monomorphic within countries. This would be expected to facilitate the derivation of SNP sets that indicate the country of origin.

#### 6.1.4 Derivation of SNP sets from merged matrices

We further demonstrated the ability of MinSNPs to analyse large datasets. To this end, we obtained additional *S. aureus* data collected through different initiatives (Bioprojects from Genbank: PRJEB3174 (20, 21), PRJEB32286 (21), and PRJNA400143 (22)) and created a large orthologous SNP matrix using a modification of the SPANDx pipeline (23) (Supplementary Information 5 (https://figshare.com/s/aadf860f3cfd9c416e3f)). The matrix encompasses 3,279 isolates (including the reference genome Mu50) and 164,335 SNP positions. We then used this matrix to validate the SNPs discriminating both ST762 and CC1 obtained earlier using only STARRS dataset. It was found that apart from one SNP set, all the previously identified single SNP sets retained 100% sensitivity and specificity for ST762 with this large data set. However, two of the SNPs were not present in the matrix. For CC1 (ST1, ST762, ST2851, ST2981), most of the previously identified SNP sets were not fully present in the matrix (i.e., the STARRS derived sets often included positions that were not included in the merged matrix due to quality filtering). For similar reasons, not all the members of previously identified high-D SNPs-sets were present in the new matrix, and no meaningful comparison between the previous analysis and current analysis could be made (see Supplementary information 5 (https://figshare.com/s/aadf860f3cfd9c416e3f)).

We also reran the same tasks in 6.1.2 with the matrix. We identified 50 individual SNPs and 50 two-member SNP sets that discriminate all ST762 isolates from all others. We similarly identified 39 individual SNPs and 61 two-member SNP sets (100 SNPs sets) that discriminate all CC1 isolates from all others.

We then experimented with the *D* mode analysis to accomplish two different tasks. First, we attempted to identify SNPs that discriminated all CCs from each other. To accomplish this, all the variant positions between isolates within the same CC were identified and recorded. A reduced matrix was then constructed that contained only a single isolate from each of the CCs. We then excluded from analysis all the previously recorded variant positions within CCs, before running a *D* mode search. It was found that a minimum of seven SNPs were required to discriminate all 33 CCs from each other. MinSNPs was tasked to provide 200 alternative SNP sets that achieved a *D* of 1.0. Of these, 165 of the sets had seven members; the remaining had eight members.

Next, we explored the resolving power of SNP sets identified simply to maximise *D*, without reference to CC. Similarly, we identified five high-*D* 10-SNP sets (Supplementary Information 5). Prior to running MinSNPs analysis, all but a subset of 100 CC22 isolates were randomly selected to be included in the input matrix to avoid overly biasing the analysis to include SNPs that discriminated within CC22. We obtained SNP sets with *D* values (recalculated using the entire matrix) ranging from 0.6314 to 0.6461. We selected the SNP set with the highest *D* value and constructed the allelic profile with the first 5 SNPs (see Supplementary Information 5 (https://figshare.com/s/aadf860f3cfd9c416e3f)). As expected from the similar experiment performed with the smaller STARRS data set, there was close but imperfect correspondence between CC and allelic profile, even though there was no reference to CC in the SNP derivation procedure (see Supplementary Information 5 for comparison).

In summary, MinSNPs provides a flexible means for deriving SNP sets from sequence alignments that are optimised for lineage-specific or generalised resolving power. We have demonstrated its utility using large data sets, where one such data set was a SNP matrix assembled from multiple *S. aureus* bioprojects, using a modified pipeline that we also report here. This provides the potential for assembling matrices encompassing the global diversity of microorganisms and mining there for optimised marker sets.

**Figure 1:**
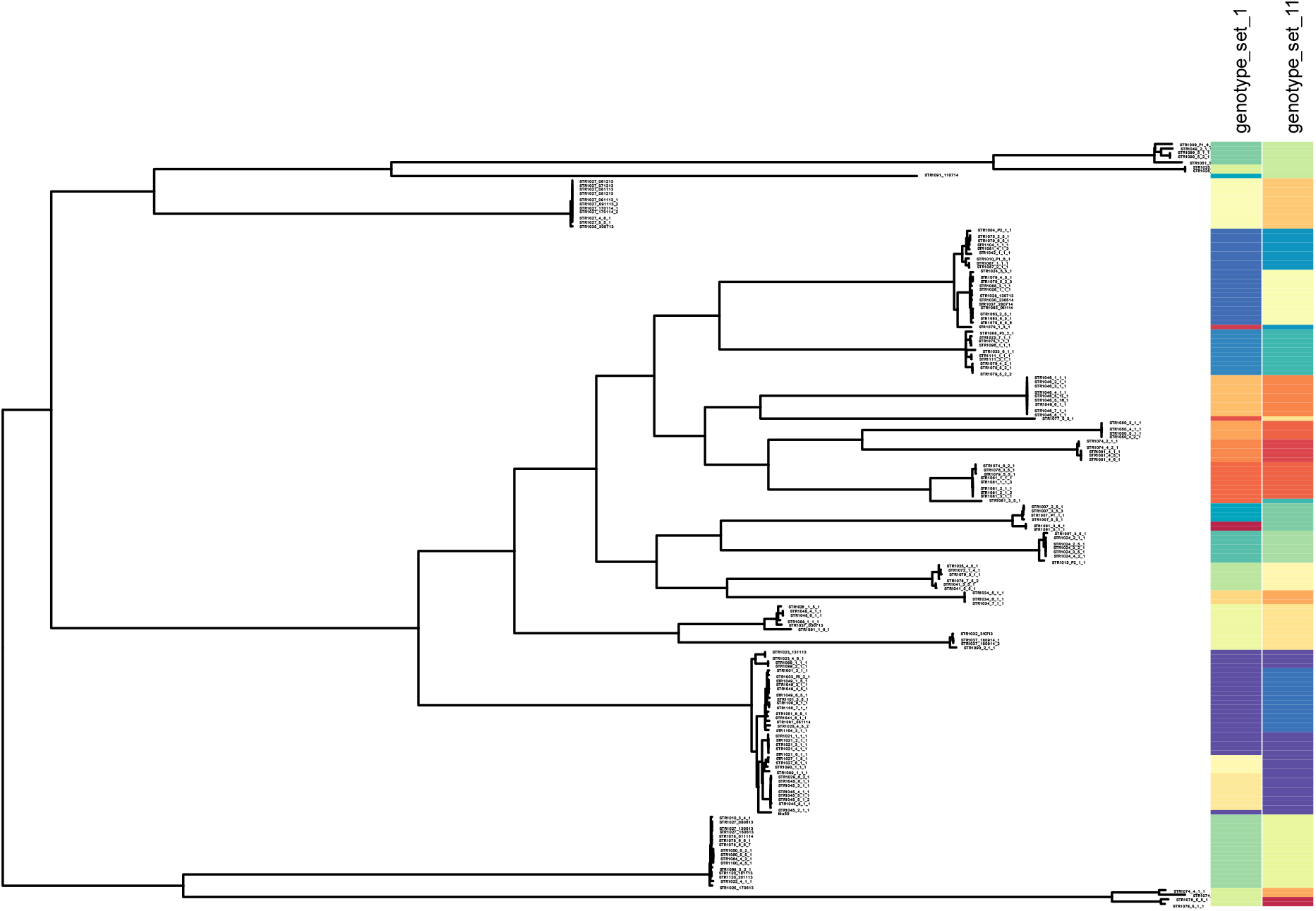
STARRS: Phylogeny and genotypes as defined by high-D SNP sets 1 and 11. The phylogenetic tree was taken from the original paper (13) and labelled with two newly identify high-*D* SNP sets. (https://microreact.org/project/minsnps-starrs). High-*D* SNP sets 1 and 11 include positions 111760, 1925985, 2663300, 2683490, 124088, and 539419, 1413096, 1146945, 2184528, 1577370 of the Mu50 reference genome respectively.

**Table 1:**
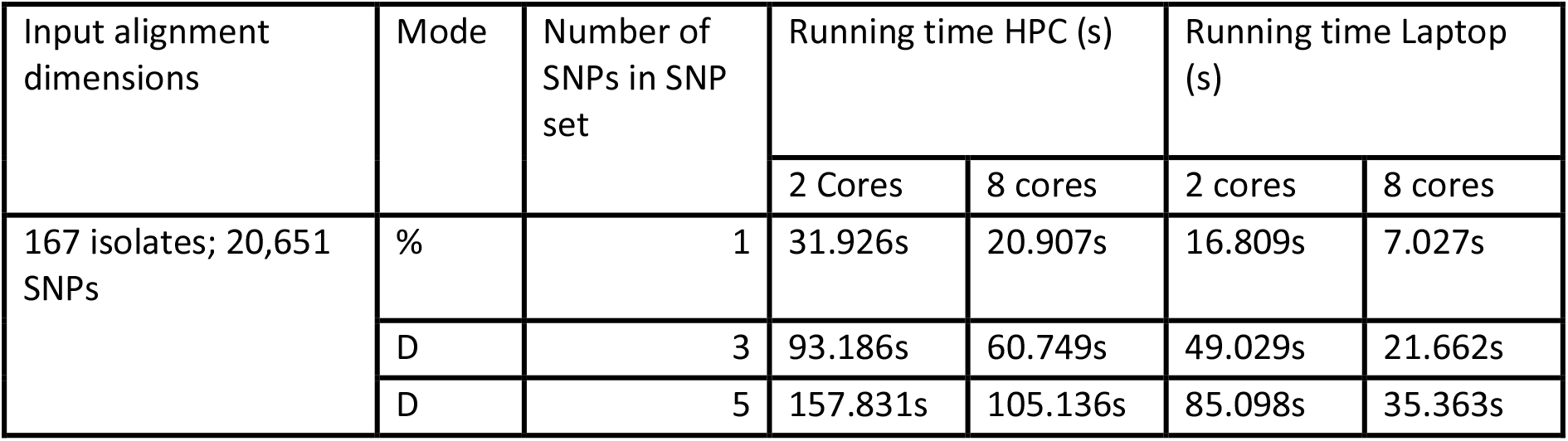
Input alignment dimensions versus run time.

| Input alignment dimensions | Mode | Number of SNPs in SNP set | Running time HPC (s) |  | Running time Laptop (s) |  |
| --- | --- | --- | --- | --- | --- | --- |
|  |  |  | 2 Cores | 8 cores | 2 cores | 8 cores |
| 167 isolates; 20,651 SNPs | % | 1 | 31.926s | 20.907s | 16.809s | 7.027s |
|  | D | 3 | 93.186s | 60.749s | 49.029s | 21.662s |
|  | D | 5 | 157.831s | 105.136s | 85.098s | 35.363s |

**Table 2:**
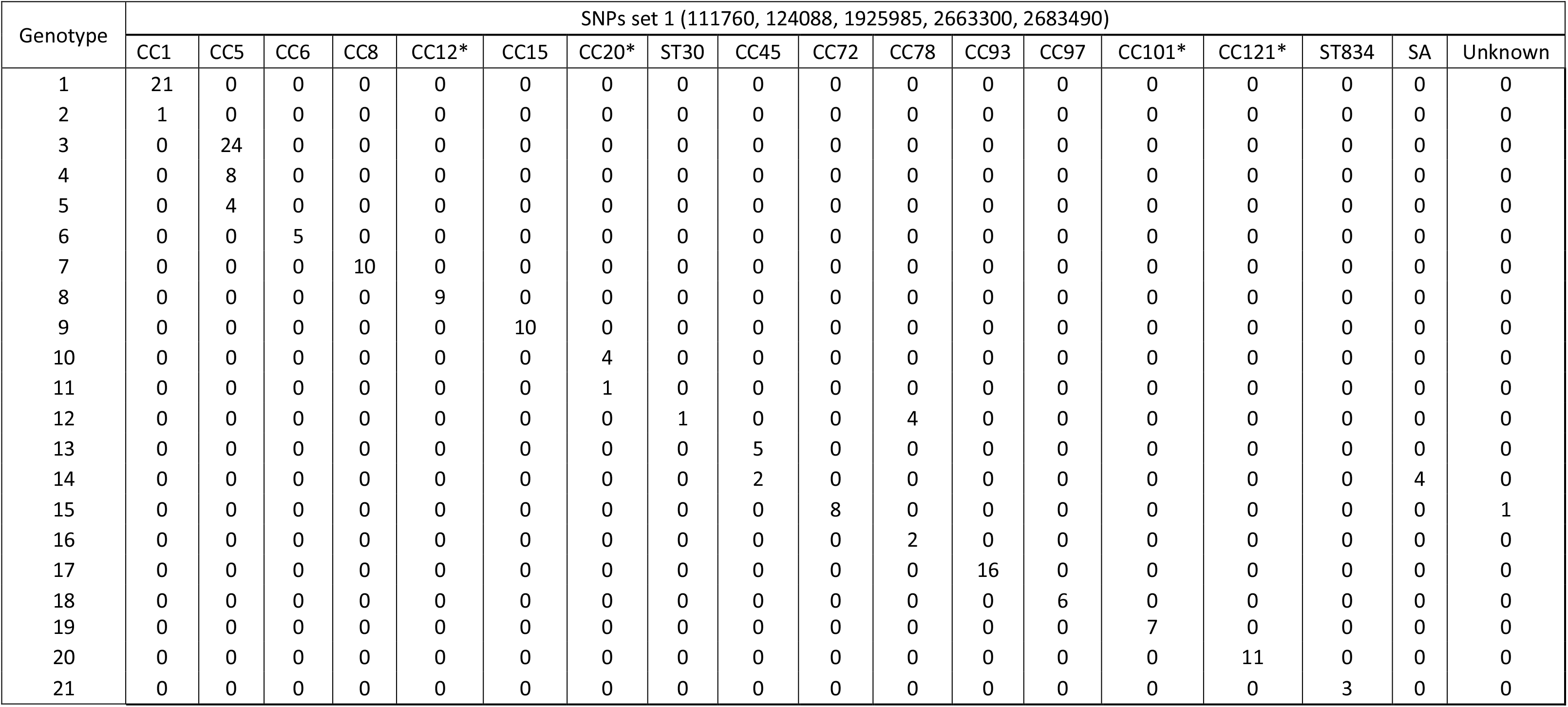
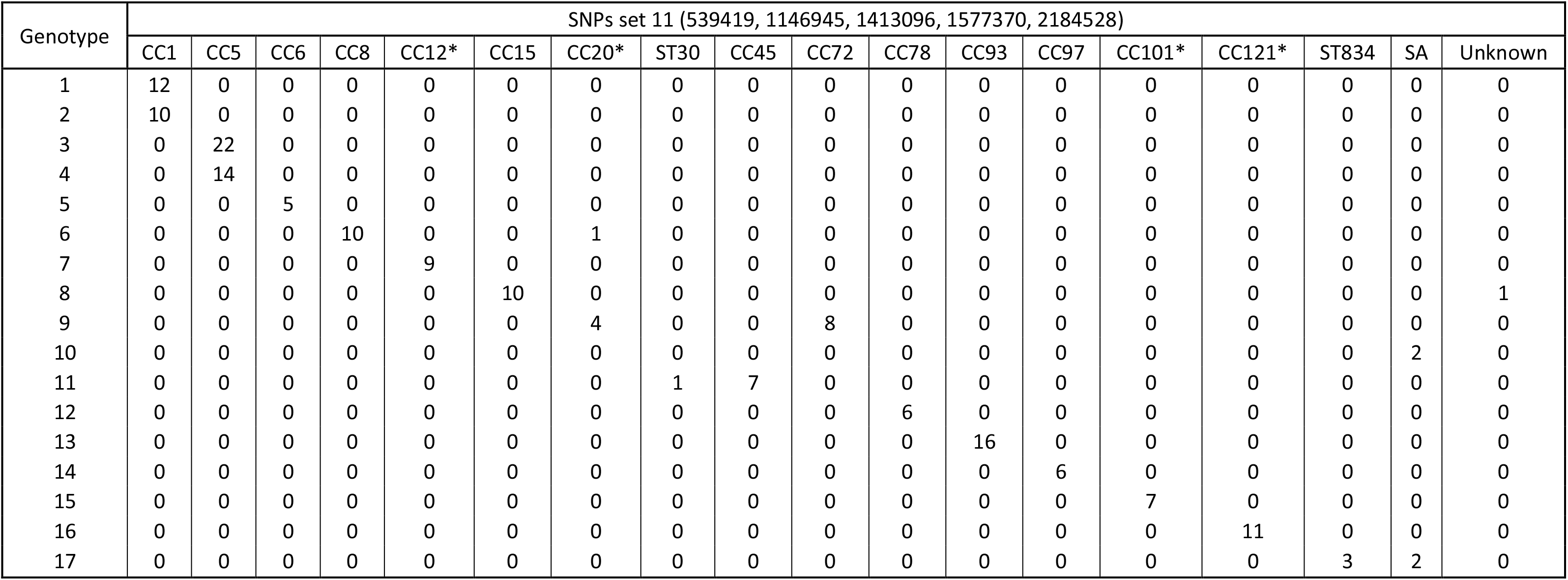
STARRS: Breakdown of CC/Singletons for genotypes defined by SNP sets 1 and 11. The distinction between singletons and CCs is somewhat arbitrary. The CCs labelled with “*” were present only as the CC founder ST in the STARRS isolates. Column SA refers to *S. argenteus*. 2a Breakdown of CC/Singletons for genotypes defined by SNPs set 1 2b Breakdown of CC/Singletons for genotypes defined by SNPs set 11

**Table 3:**
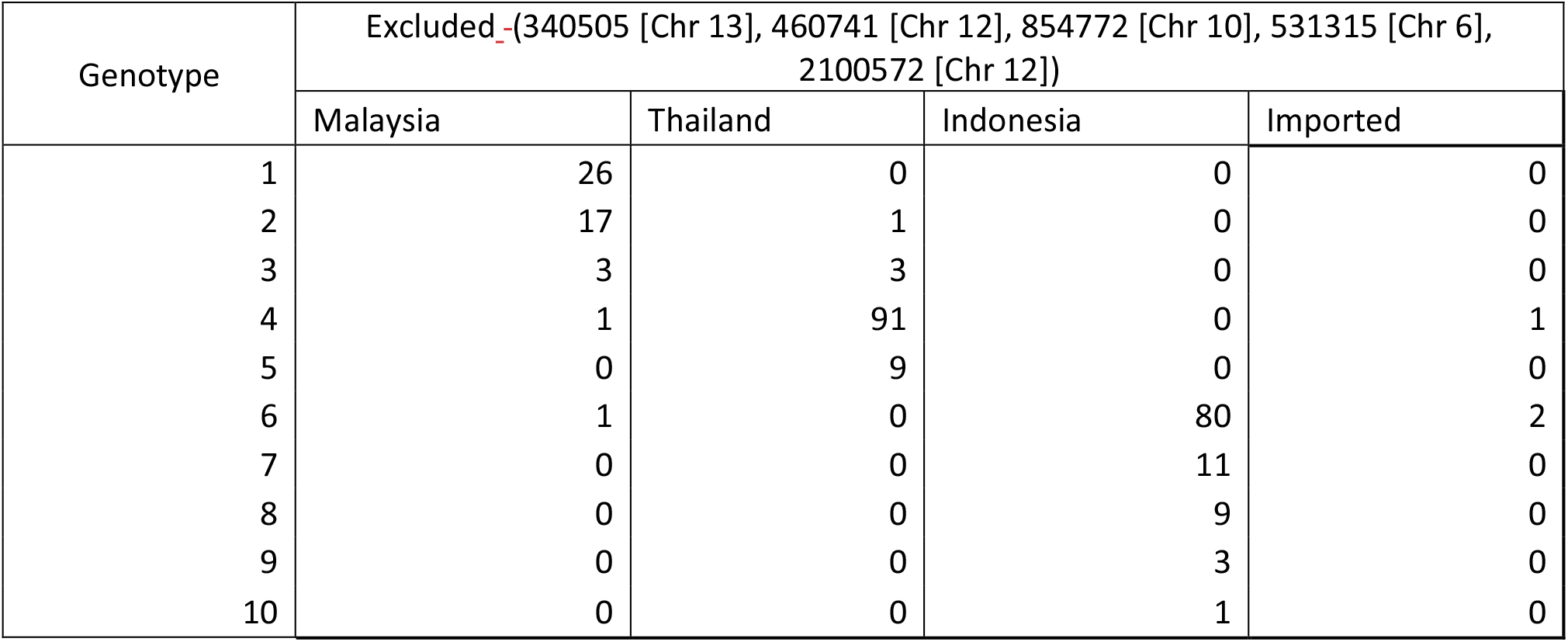
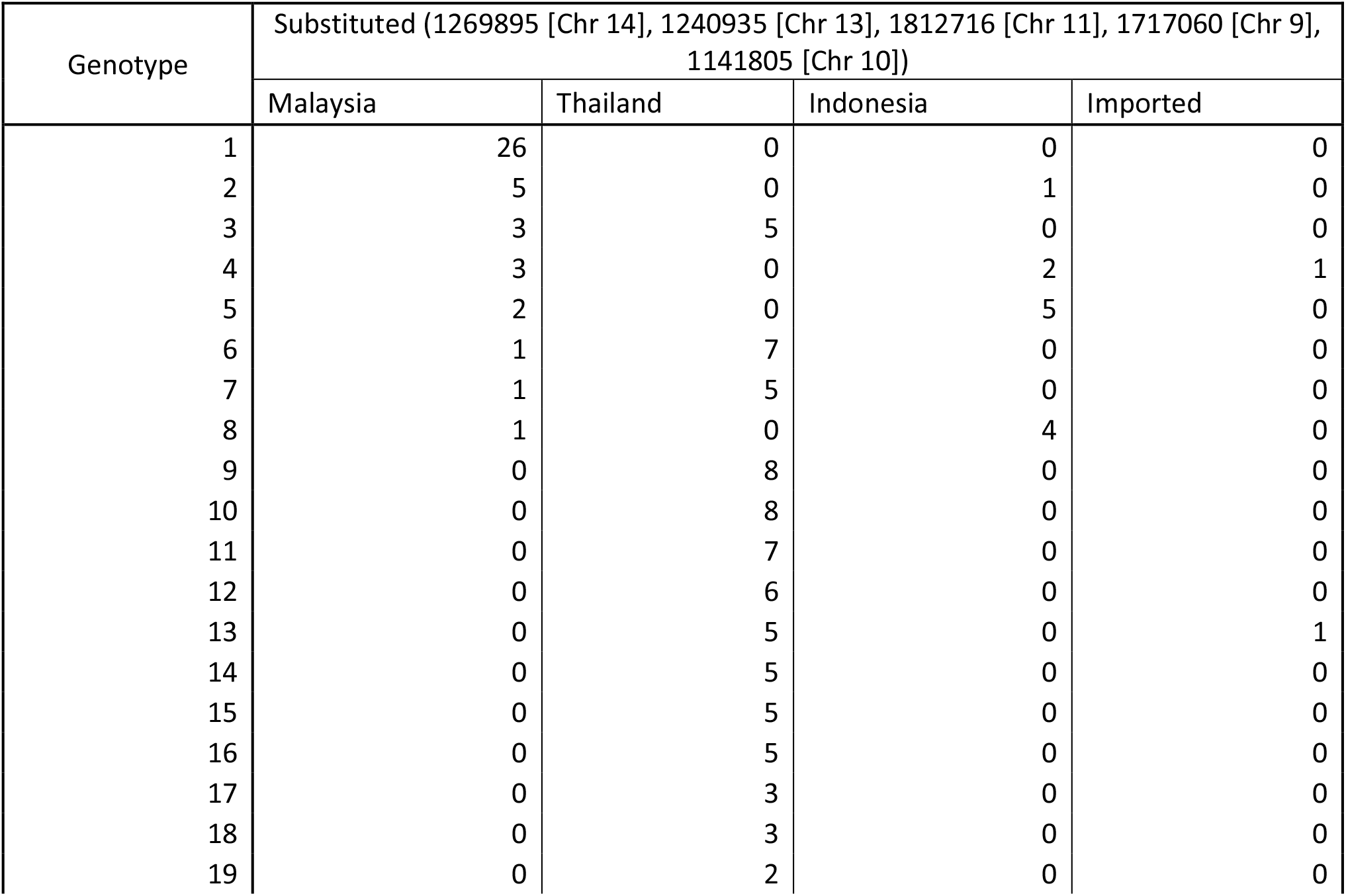
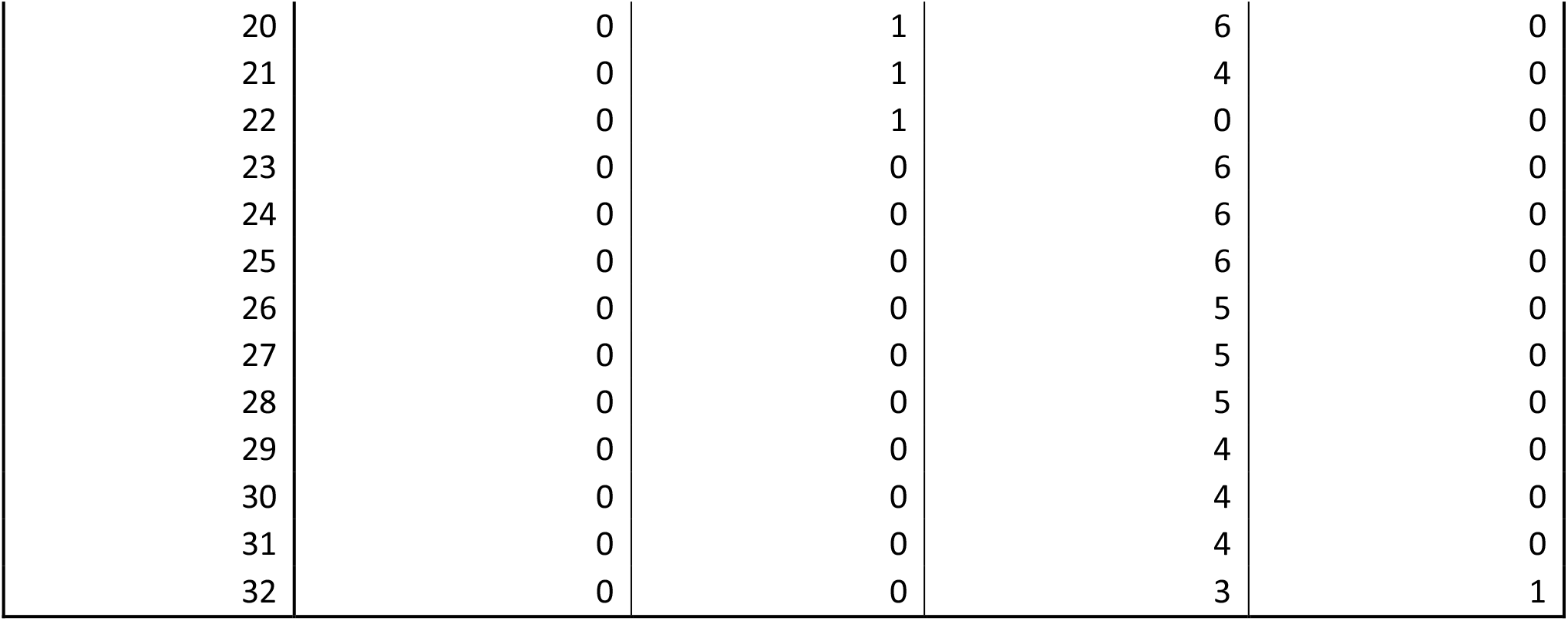
*P. vivax*: Genotypes defined by high-*D* SNP set 1 (ambiguity codes excluded vs substituted) 1a Genotypes defined by high-D SNPs set 1 (ambiguity code excluded) 3b Genotypes defined by high-D SNPs set 1 (ambiguity code substituted)

| Genotype | Excluded (340505 [Chr 13], 460741 [Chr 12], 854772 [Chr 10], 531315 [Chr 6], 2100572 [Chr 12]) |  |  |  |
| --- | --- | --- | --- | --- |
|  | Malaysia | Thailand | Indonesia | Imported |
| 1 | 26 | 0 | 0 | 0 |
| 2 | 17 | 1 | 0 | 0 |
| 3 | 3 | 3 | 0 | 0 |
| 4 | 1 | 91 | 0 | 1 |
| 5 | 0 | 9 | 0 | 0 |
| 6 | 1 | 0 | 80 | 2 |
| 7 | 0 | 0 | 11 | 0 |
| 8 | 0 | 0 | 9 | 0 |
| 9 | 0 | 0 | 3 | 0 |
| 10 | 0 | 0 | 1 | 0 |

| Genotype | Substituted (1269895 [Chr 14], 1240935 [Chr 13], 1812716 [Chr 11], 1717060 [Chr 9], 1141805 [Chr 10]) |  |  |  |
| --- | --- | --- | --- | --- |
|  | Malaysia | Thailand | Indonesia | Imported |
| 1 | 26 | 0 | 0 | 0 |
| 2 | 5 | 0 | 1 | 0 |
| 3 | 3 | 5 | 0 | 0 |
| 4 | 3 | 0 | 2 | 1 |
| 5 | 2 | 0 | 5 | 0 |
| 6 | 1 | 7 | 0 | 0 |
| 7 | 1 | 5 | 0 | 0 |
| 8 | 1 | 0 | 4 | 0 |
| 9 | 0 | 8 | 0 | 0 |
| 10 | 0 | 8 | 0 | 0 |
| 11 | 0 | 7 | 0 | 0 |
| 12 | 0 | 6 | 0 | 0 |
| 13 | 0 | 5 | 0 | 1 |
| 14 | 0 | 5 | 0 | 0 |
| 15 | 0 | 5 | 0 | 0 |
| 16 | 0 | 5 | 0 | 0 |
| 17 | 0 | 3 | 0 | 0 |
| 18 | 0 | 3 | 0 | 0 |
| 19 | 0 | 2 | 0 | 0 |
| 20 | 0 | 1 | 6 | 0 |
| 21 | 0 | 1 | 4 | 0 |
| 22 | 0 | 1 | 0 | 0 |
| 23 | 0 | 0 | 6 | 0 |
| 24 | 0 | 0 | 6 | 0 |
| 25 | 0 | 0 | 6 | 0 |
| 26 | 0 | 0 | 5 | 0 |
| 27 | 0 | 0 | 5 | 0 |
| 28 | 0 | 0 | 5 | 0 |
| 29 | 0 | 0 | 4 | 0 |
| 30 | 0 | 0 | 4 | 0 |
| 31 | 0 | 0 | 4 | 0 |
| 32 | 0 | 0 | 3 | 1 |

## 7. Author statements

### 7.1 Authors and contributors

KSH: Data curation, Formal analysis, Investigation, Methodology, Software, Visualisation, Writing-original draft, Writing – review and editing.

DCH: Data curation, Methodology, Resources, Project Administration, Supervision, Writing-review and editing.

SA: Methodology, Resources, Supervision, Writing-review and editing.

PS: Funding acquisition, Methodology, Software, Supervision, Writing-Review and Editing

PMG: Conceptualisation, Formal analysis, Funding acquisition, Investigation, Methodology, Project Administration, Resources, Supervision, Writing – original draft, Writing-review and editing.

### 7.4 Ethical approval

This work does not involve human or animal research. The research team received written confirmation of this from the Human Research Ethics Committee for the Northern Territory Government Department of Health and the Menzies School of Health research.

### 7.5 Consent for publication

Not applicable

## 7.6 Acknowledgements

The authors thank Mariana Barnes and Tegan Harris from the Menzies School of Health Research for assistance with the installation of MinSNPs onto the Charles Darwin University high performance computer cluster, and also with the associated software documentation tasks.

## References

1. Kim T-W, Jang Y-H, Jeong MK, Seo Y, Park CH, Kang S, et al. Single-nucleotide polymorphism-based epidemiological analysis of Korean Mycobacterium bovis isolates. Journal of Veterinary Science. 2021;22(2):e24–e.

2. Napier G, Campino S, Merid Y, Abebe M, Woldeamanuel Y, Aseffa A, et al. Robust barcoding and identification of Mycobacterium tuberculosis lineages for epidemiological and clinical studies. Genome Medicine. 2020;12(1):114–.

3. Vorimore F, Aaziz R, de Barbeyrac B, Peuchant O, Szymańska-Czerwińska M, Herrmann B, et al. A New SNP-Based Genotyping Method for C. psittaci: Application to Field Samples for Quick Identification. Microorganisms. 2021;9(3):625–.

4. Jehanne Q, Pascoe B, Bénéjat L, Ducournau A, Buissonnière A, Mourkas E, et al. Genome-Wide Identification of Host-Segregating Single-Nucleotide Polymorphisms for Source Attribution of Clinical Campylobacter coli Isolates. Applied and Environmental Microbiology. 2020;86(24):e01787–20–e–20.

5. Kato CY, Chung IH, Robinson LK, Eremeeva ME, Dasch GA. Genetic typing of isolates of Rickettsia typhi. PLoS Neglected Tropical Diseases. 2022;16(5):e0010354–e.

6. Rahman M-M, Lim S-J, Park Y-C. Development of Single Nucleotide Polymorphism (SNP)-Based Triplex PCR Marker for Serotype-specific Escherichia coli Detection. Pathogens. 2022;11(2):115–.

7. Jacob CG, Thuy-Nhien N, Mayxay M, Maude RJ, Quang HH, Hongvanthong B, et al. Genetic surveillance in the Greater Mekong subregion and South Asia to support malaria control and elimination. eLife. 2021;10:e62997–e.

8. Robertson GA, Thiruvenkataswamy V, Shilling H, Price EP, Huygens F, Henskens FA, et al. Identification and interrogation of highly informative single nucleotide polymorphism sets defined by bacterial multilocus sequence typing databases. J Med Microbiol. 2004;53(Pt 1):35–45.

9. Price EP, Inman-Bamber J, Thiruvenkataswamy V, Huygens F, Giffard PM. Computer-aided identification of polymorphism sets diagnostic for groups of bacterial and viral genetic variants. BMC Bioinformatics. 2007;8:278.

10. Tong SYC, Xie S, Richardson LJ, Ballard SA, Dakh F, Grabsch EA, et al. High-Resolution Melting Genotyping of Enterococcus faecium Based on Multilocus Sequence Typing Derived Single Nucleotide Polymorphisms. PLOS ONE. 2011;6(12):e29189–e.

11. Price EP, Inman-Bamber J, Thiruvenkataswamy V, Huygens F, Giffard PM. Computer-aided identification of polymorphism sets diagnostic for groups of bacterial and viral genetic variants. BMC Bioinformatics. 2007;8(1):278–.

12. Giffard PM, Andersson P, Wilson J, Buckley C, Lilliebridge R, Harris TM, et al. CtGEM typing: Discrimination of Chlamydia trachomatis ocular and urogenital strains and major evolutionary lineages by high resolution melting analysis of two amplified DNA fragments. PLOS ONE. 2018;13(4):e0195454–e.

13. Holt DC, Harris TM, Hughes JT, Lilliebridge R, Croker D, Graham S, et al. Longitudinal whole-genome based comparison of carriage and infection associated Staphylococcus aureus in northern Australian dialysis clinics. 2021;16(2):e0245790–e.

14. Lilliebridge RA, Tong SY, Giffard PM, Holt DC. MLST based Staphylococcus aureus typing scheme using high-resolution melting analysis of SNP nucleated PCR fragments. The clinical and molecular epidemiology of community-associated Staphylococcus aureus in northern Australia. 2010:119–.

15. Robertson G, Thiruvenkataswamy V, Shilling H, Price E, Huygens F, Henskens F, et al. Identification and Interrogation of Highly Informative Single Nuceotide Polymorphism Sets Defined by Bacterial Multilocus Sequence Typing Databases. Journal of medical microbiology. 2004;53:35–45.

16. Noviyanti R, Miotto O, Barry A, Marfurt J, Siegel S, Thuy-Nhien N, et al. Implementing parasite genotyping into national surveillance frameworks: feedback from control programmes and researchers in the Asia–Pacific region. BioMed Central; 2020.

17. Fola AA, Kattenberg E, Razook Z, Lautu-Gumal D, Lee S, Mehra S, et al. SNP barcodes provide higher resolution than microsatellite markers to measure Plasmodium vivax population genetics. Malaria Journal. 2020;19:375–.

18. Diez Benavente E, Campos M, Phelan J, Nolder D, Dombrowski JG, Marinho CRF, et al. A molecular barcode to inform the geographical origin and transmission dynamics of Plasmodium vivax malaria. PLoS Genetics. 2020;16(2):e1008576–e.

19. Auburn S, Benavente ED, Miotto O, Pearson RD, Amato R, Grigg MJ, et al. Genomic analysis of a pre-elimination Malaysian Plasmodium vivax population reveals selective pressures and changing transmission dynamics. Nature Communications. 2018;9(1):2585–.

20. Toleman MS, Reuter S, Coll F, Harrison EM, Blane B, Brown NM, et al. Systematic Surveillance Detects Multiple Silent Introductions and Household Transmission of Methicillin-Resistant Staphylococcus aureus USA300 in the East of England. The Journal of Infectious Diseases. 2016;214(3):447–53-–53.

21. Coll F, Raven KE, Knight GM, Blane B, Harrison EM, Leek D, et al. Definition of a genetic relatedness cutoff to exclude recent transmission of meticillin-resistant Staphylococcus aureus: a genomic epidemiology analysis. The Lancet Microbe. 2020;1(8):e328–e35–e–e35.

22. Manara S, Pasolli E, Dolce D, Ravenni N, Campana S, Armanini F, et al. Whole-genome epidemiology, characterisation, and phylogenetic reconstruction of Staphylococcus aureus strains in a paediatric hospital. Genome medicine. 2018;10(1):1–19–1–.

23. Sarovich DS, Price EP. SPANDx: a genomics pipeline for comparative analysis of large haploid whole genome re-sequencing datasets. BMC research notes. 2014;7(1):1–9–1–9.

